# Novel Quorum Sensing Activity in East Antarctic Soil Bacteria

**DOI:** 10.1101/749861

**Authors:** Sin Yin Wong, James C. Charlesworth, Nicole Benaud, Brendan P. Burns, Belinda C. Ferrari

**Author notes:** Author contributions: B.C.F, B.P.B, J.C.C and S.Y.W designed the study. N.B performed genome sequence analyses. S.Y.W performed the experiments, analysed the data and drafted the manuscript. All authors edited the final manuscript.

## Abstract

Antarctica, being the coldest, driest and windiest continent on Earth, represents the most extreme environment a living organism can thrive in. Under constant exposure to harsh environmental threats, terrestrial Antarctica remains home to a great diversity of microorganisms, indicating that the soil bacteria must have adapted a range of survival strategies that require cell-to-cell communication. Survival strategies include secondary metabolite production, biofilm formation, bioluminescence, symbiosis, conjugation, sporulation and motility, all of which are often regulated by quorum sensing (QS), a type of bacterial communication. Up to now, such mechanisms have not been explored in terrestrial Antarctica. Here, for the first time, LuxI/LuxR-based quorum sensing (QS) activity was delineated in soil bacterial isolates recovered from Adams Flat, in the Vestfold Hills region of East Antarctica. Interestingly, we identified the production of potential homoserine lactones (HSLs) ranging from medium to long chain length in 19 bacterial species using three biosensors, namely *Agrobacterium tumefaciens* NTL4, *Chromobacterium violaceum* CV026 and *Escherichia coli* MT102, in conjunction with thin layer chromatography (TLC). The majority of detectable HSLs were from gram-positive microorganisms not previously known to produce HSLs. This discovery further expands our understand of the microbial community capable of this type of communication, as well as providing insights into physiological adaptations of microorganisms that allow them to survive in the harsh Antarctic environment.

**IMPORTANCE:** Quorum sensing, a type of bacterial communication, is widely known to regulate many processes including those that confer survival advantage. However, little is known about communication by bacteria thriving within Antarctic soils. Employing a combination of bacteria biosensors, analytical techniques, and genome mining, we found a variety of Antarctic soil bacteria speaking a common language, via the LuxI/LuxR-based quorum sensing, thus potentially supporting survival in a mixed microbial community. This is the first report of quorum sensing activity in Antarctic soils and has provided a platform for studying physiological adaptations of microorganisms that allow them to not just survive but thrive in the harsh Antarctic environment.

## INTRODUCTION

Terrestrial Antarctica represents one of the most extreme environments a living organism can thrive in (1, 2). Precipitation is limited (3), and of this, the majority falls as snow and ice crystals that do not melt but build up over time to form massive ice sheets. Cold, dense air from these elevated ice-built surfaces continuously sinks downhill and is replaced by subsiding air from above, resulting in strong winds that increases the aridity of Antarctica. The high elevation together with reflection of sunlight by the snow and ice cover, not allowing heat energy absorption, contribute to the low temperatures reaching −89 °C once in 1983 (4, 5). Antarctica experiences total darkness during winter, while during summer, it receives 24-hour-sunlight, and with this, damaging UVB rays (1, 6). The harsh environmental conditions of Antarctica have consequently shaped the simple trophic structures present in Antarctic soil ecosystems (7). The lack of vascular plants (except for the Antarctic Peninsula), the small proportion of macroinvertebrates and the low water availability, leads to extremely low organic matter content (8, 9).

Despite the harsh conditions, terrestrial Antarctica remains home to a great diversity of microorganisms, ranging from phyla frequently observed in soils from temperate regions, such as Proteobacteria, Actinobacteria, Bacteroidetes and Firmicutes (2, 8, 10) through to Candidate phyla which have no cultured representatives (9, 11, 12). From an ecological standpoint, these Antarctic microorganisms may have evolved under the environmental stressors such as high UV exposure during summer, complete absence of light during winter, frequent freeze-thaw cycles, osmotic limitation and nutrient starvation, to possess unique biochemical adaptations that confer them a selective advantage in the seemingly inhospitable terrain (9-11, 13, 14). These adaptations are likely to incorporate a variety of survival strategies such as secondary metabolite production (15–18), biofilm formation (19–21), bioluminescence, symbiosis, conjugation, sporulation and motility, all of which are often regulated by quorum sensing (QS), a type of bacterial communication (22–24).

The most widely reported QS system is the LuxI/LuxR system (25, 26) that utilizes homoserine lactones (HSLs) as signal molecules. HSLs are a diverse group of molecules synthesized by the LuxI synthase enzyme. The typical structure of HSL consists of a conserved homoserine lactone ring and a side chain of variable length that can undergo various types of modification such as substitution, hydroxylation and carboxylation at the C-3 position, allowing the signalling molecules to be species-specific (27). The side chain of HSLs is usually a fatty acyl group but new classes of HSLs containing p-coumaryl group (28), cinnamoyl group (29) and aryl group (30, 31) have also been found to be produced and detected by LuxI/LuxR homologs. In addition, LuxR were found to be capable of binding non-HSL signalling molecules derived from other organisms, such as cyclic diguanylate (c-di-GMP), furanones, and diketopiperazines (DKPs) (32), overcoming communication barriers between species, or even kingdoms.

While the LuxI/LuxR system was previously thought to be exclusive to gram-negative bacteria, knowledge on the distribution of this QS system among microbes is growing, with HSL activity now reported in Archaea (33–35), in gram-positive marine bacteria (36, 37), gram-positive rhizobacteria from a mangrove swamp (38) and gram-positive bacteria from hypersaline microbial mats (39). In Antarctica, a limited number of HSL-based QS studies is being restricted to marine environments (24, 40). The occurrence of HSL activity in Antarctic soil bacteria is not known, thus major gaps remain in the understanding of the role that quorum sensing may play in nutrient-poor, cold desert environments (27).

Here, we investigated the potential of HSL-based quorum sensing by Antarctic soil bacteria, specifically from a hyper-arid site named Adams Flat, in the Vestfold Hills region. Adams Flat soils are comprised of a microbial diversity with dominance by the phylum Actinobacteria (41) (Supplementary Figure 1), combined with a high proportion of bacteria with the genetic capacity for diverse secondary metabolite production (15). Microbial communities in Adams Flat were also recently shown to be carrying out significant levels of atmospheric chemosynthesis to support primary production (41), hence suggesting the need to evolve with novel functionalities in order to survive in a barren desert environment.

## MATERIAL & METHODS

### Sample Site and Description

All soil samples were collected by the Australian Antarctic Division (AAD), between 2005 and 2012, and were stored at −80 °C until further analysis (9, 42). This includes nine sites across East Antarctica spanning two different regions, the Windmill Islands and Vestfold Hills (Supplementary Figure 1). Soils from Vestfold Hills are hyper-arid as compared to arid soils in the Windmill Islands. This means Vestfold Hills have lower levels of organic carbon, nitrogen and moisture (9, 41, 43). As no cultivation studies have been performed for Adams Flat (68°33’S, 78°1’E), three soil samples, (designated as Soil 1, 2 and 3 in this paper) that exhibited a diversity in measured soil factors, of organic chemical contents and physical parameters including pH and moisture were selected for cultivation (Supplementary Table 1).

### Cultivation onto Standard Media

Soil inocula were prepared from the three Adams Flat soil samples by adding 1 g soil to 5 ml 0.9% sterile saline. The soil suspension was serially diluted and plated onto ¾ nutrient agar (Oxoid^TM^). Culture plates were left to incubate at RT (22 °C) for 30 days. Pure cultures were established for each morphologically different isolate by successive sub-culturing onto ¾ nutrient agar (Oxoid^TM^). All isolates were preserved in cryovials (Microbank^TM^, Pro Lab Diagnostics) at −80 °C for future use.

### DNA extraction and 16S rDNA Gene Sequencing of Isolates

DNA extractions were performed using an in-house modified bead beating protocol based on the FastPrep® homogenisation system (MP Biomedicals, California). A single colony was transferred into a 2 ml microcentrifuge tube containing 600 μl autoclaved Milli-Q® water and 0.5 g of both the 0.1 mm and 0.5 mm diameter glass beads (MoBio Inc, Carlsbard). The mixture was homogenized in FastPrep®120 cell disrupter, at speed setting 6.0 for 40 sec. Following incubation at 95 °C for 5 min, the sample was centrifuged at 20800 x g for 3 min and 400 μl of the supernatant was transferred to a new tube. This crude DNA extract was then purified via an ethanol purification protocol as described in (44). The primer set 27F and 1492R (45) was used to amplify the bacterial conserved 16S rDNA genes in the extracted DNA samples prior to sequencing via the Sanger method at the Ramaciotti Centre for Gene Function Analysis (UNSW Sydney, Sydney). Obtained sequences were trimmed before comparisons with reference sequences in the National Centre for Biotechnology Information (NCBI) nucleotide database using BLASTn search tool.

### Preliminary Screening for HSL-Producing Strains

The bioreporter strains used were *C. violaceum* CV026 (detects C4-, C6-, C8-, 3-oxo-C6- and 3-oxo-C8-HSLs) (46), *A. tumefaciens* NTL4 (pZLR4) (detects C6 - C14-HSLs, all 3-oxo-HSLs, and 3-hydroxy-HSLs of C6 – C10) (47) and *E. coli* MT102 (pjBA132) (detects C4 – C12-HSLs, 3-oxo-HSLS of C6, C8 and C10) (48). The positive control used was *Pseudomonas Aeruginosa* PAOI. All the above strains were kindly gifted by Dr. Onder Kimyon (School of Civil and Environmental Engineering, UNSW, Sydney). Antibiotics (Sigma-Aldrich) were used at the following concentrations: for *E. coli* MT102, Ampicillin (100 µg/ml); for *A. tumefaciens* NTL4, Spectinomycin (50 µg/ml), and Tetracycline (4.5 µg/ml); for *C. violaceum* CV026, Kanamycin (20 µg/ml).

HSL activity of bacterial isolates was tested using the biosensors *C. violaceum* CV026 and *A. tumefaciens* NTL4 via the cross-feeding plate assay as described in (49), and by *E. coli* MT102 biosensor via the well-plate assay as in (39). Briefly, each test isolate was streaked side to side with the biosensors CV026 and NTL4 on ¾ nutrient agar supplemented with 40 µg /ml X-gal (Invitrogen). Self-streaks of each biosensors served as negative controls and a streak of *P. aeruginosa* PAOI with the biosensors served as a positive control. These cross-feeding plates were incubated at RT for a week and were monitored for positive HSL activity as indicated by visible purple or blue pigmentation, respectively.

For the bioassay against *E. coli* MT102, test isolates were grown in 25 ml of ¾ nutrient broth (Oxoid^TM^) cultures at RT while shaking at 100 rpm until stationary phase, at an approximate OD_600_ of 0.5 – 1.0. Absorbance at 600 nm was measured right after inoculation (T= 0) and subsequently after every 24 h for 7 d. Cell cultures were subjected to ethyl acetate extraction, followed by incubation with *E. coli* MT102, and analysis as per (39).

### Thin layer chromatography profiling of presumptive HSLs

Isolates that returned positive in the preliminary screening were grown in larger batches of culture (450 ml) to extract higher concentrations of HSLs during stationary phase, using the aforementioned ethyl acetate extraction method. Growth curves were determined for all non-sporulating isolates by measuring absorbance at 600 nm starting from inoculation point and subsequently every 24 h for 3 weeks.

Culture extracts of 4-40 µl, depending of the concentration of the extracts, were spotted onto non-fluorescent reverse-phase C18 TLC glass plates coated with a silica gel matrix (Analtech TLC uniplates, Sigma-Aldrich). The chromatogram was developed with a HPLC-grade methanol and Milli-Q water mixture (60:40 v/v) as described by (50). After development, the solvent was evaporated before being overlaid with the biosensor *E. coli* MT102. A 5 ml overnight culture of *E.coli* MT102 was inoculated in 75 ml of LB broth and grown to late exponential phase by incubation at 30 °C with shaking at 150 rpm for 4 h. The entire 80 ml of culture was added to 120 ml of agar-enriched LB medium to give a final agar concentration of 0.8% (w/v) before pouring over the dried chromatogram. The TLC plates were incubated O/N at 30 °C before visualization of the active compounds using a GE phosphorimager (Fujifilm LPB filter set, 473 nm laser) at an 8-h-and 18-h time point.

Chain lengths of the presumptive HSLs were roughly estimated by comparing the *Rf* values and the shape of spots with a range of N-acyl-_L_-homoserine lactones, including N-butyryl (C4)-, N-hexanoyl (C6)-, N-heptanoyl (C7)-, N-octanoyl (C8)-, N-decanoyl (C10)-, N-dodecanoyl (C12)-HSLs, N-(3-oxo-hexanoyl)-_L_-HSL (OHHL), N-(3-oxo-octanoyl) _L_-HSL (OOHL), N-(3-oxo-decanoyl) _L_-HSL (ODHL) and N-(3-oxo-dodecanoyl) _L_-HSL (OdDHL) (Sigma-Aldrich).

### Bacterial genome sequencing and assembly for eight Antarctic isolates

In addition to Adams Flat bacterial isolates recovered in this study, eight isolates from the collection of bacteria recovered from East Antarctic Windmills Islands region (Supplementary Table 1) were included in the HSL screening assays due to availability of full genome sequences (N. Benaud, Unpublished data). Genomic DNA was sequenced using the PacBio RS II instrument with P6/C4 chemistry, and *de novo* assembly performed using FALCON v 1.8.6 (51). Assemblies were joined and circularized using Circlator v1.4.0 (52), then subjected to two rounds of consensus polishing via the Arrow algorithm of the GenomicConsensus package tool variantCaller v.2.2.1 (51). Annotations were performed on contigs using Prokka v1.13 (53). Protein sequences were phylogenetically classified into clusters of orthologous groups (COGs), assigned via the COG functional annotator module of WebMGA (http://weizhong-lab.ucsd.edu/webMGA/) (54, 55). Genome assemblies were quality assessed for completeness and contamination using CheckM v1.0.7 (56).

### Genome mining for LuxI and LuxR homologs

Potential LuxI- and LuxR-containing sequences within the eight genome sequences of Windmill Isolates were extracted from InterproScan v5.25-64.0 (57). InterProScan integrates data from various major protein signature databases: CATH-Gene3D, CDD, HAMAP, PANTHER, Pfam, PIRSF, PRINTS, ProDomBlast, PROSITE (Patters and profiles), SFLD, SMART, SUPERFAMILY and TIGRFAMs. These genomes were searched for conserved domains within autoinducer synthases and LuxR-type transcriptional regulators (Supplementary Table 3). The genomes were also searched for acyl-homoserine lactone (AHL) synthase COG category COG3916.

Ligand-binding site of the potential LuxR homologs extracted from genomes were performed with COACH software (58, 59). Following structural prediction of the receptor via the I-TASSER suite, potential ligands were determined based on structure of the binding sites via five methods (TM-SITE, S-SITE, COFACTOR, FINDSITE and ConCavity). These potential LuxI and LuxR were aligned against *V. fishcherii* LuxI (P12747) and LuxR (P12746), respectively using Clustal Omega (60). It employs an mBED algorithm to calculate seeded guide trees and HMM profile-profile techniques to generate alignments between sequences.

Phylogenetic trees were built via maximum-likelihood method based on 1000 bootstrap replications, via PhyML (61). Reference amino acid sequences were retrieved from UniProtKB database (62). Sequences to be passed through PhyML were first aligned using MUSCLE (63) in full processing mode.

## RESULTS & DISCUSSION

### Culturable diversity of Adams Flat soils

In total, 81 isolates were cultured that represented 21 different bacterial strains (Table 1). These 21 strains, belonging to 18 species, spanned four soil phyla (Figure 1), with the majority being members of Actinobacteria (58%), followed by Firmicutes (31%), Bacteroidetes (7%) and Proteobacteria (4%), which are the four most well-characterised phyla as they are those most readily culturable (2, 64). These phyla are ubiquitous and common in desert environments, including Antarctic soils (8, 20). The bacterial 16S rDNA sequences have been submitted to GenBank Database and the accession number is MN365277-MN365297.

**Figure 1.**
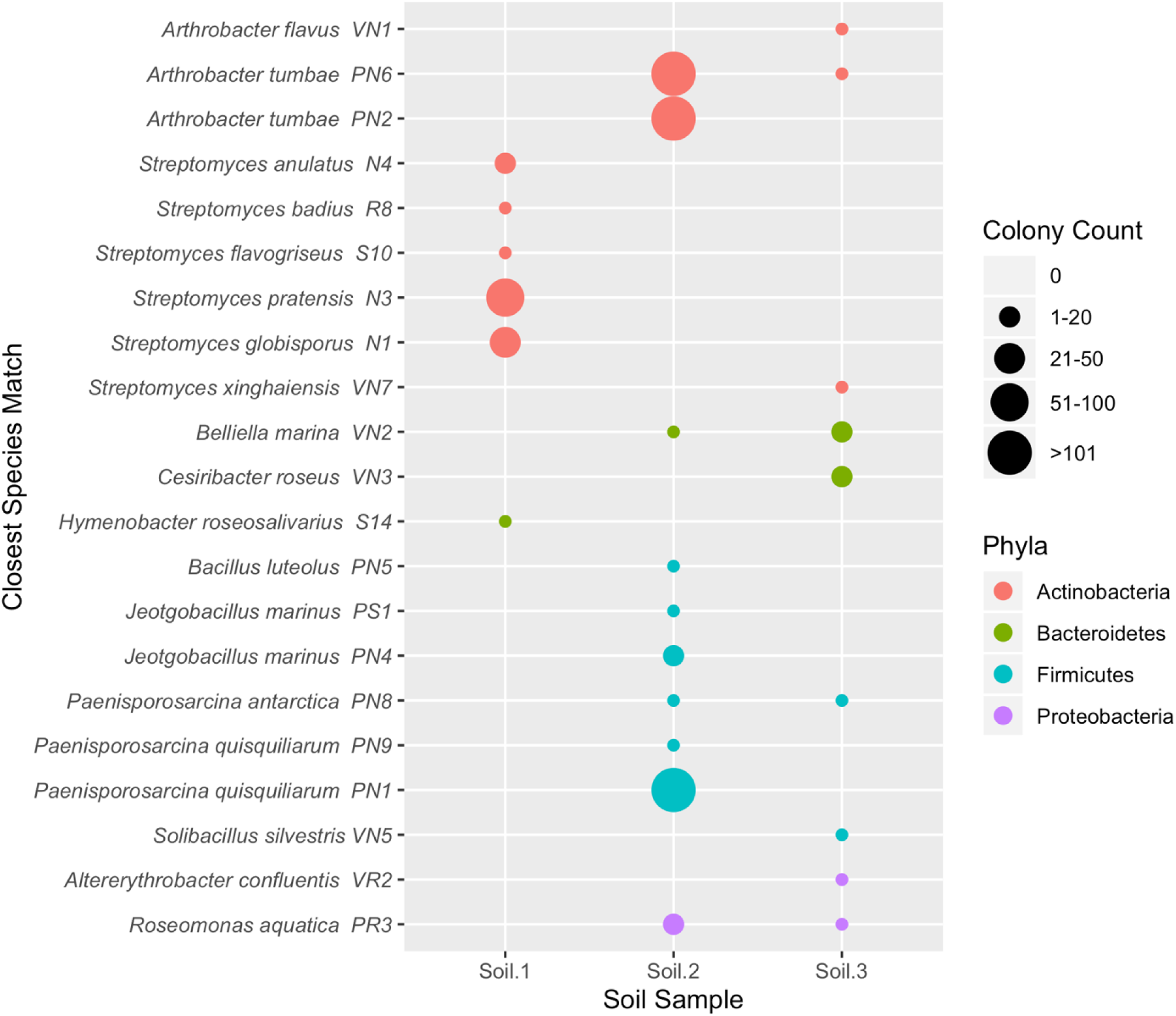
Presumptive bacterial species cultured from all three Adams Flat soil samples. Soil 2 and Soil 3, with higher total organic content compared to Soil 1, recovered the highest diversity of taxa. As expected, Actinobacteria dominated the culturable fraction, especially in Soil 1 and Soil 2.

**Table 1.** Identification of isolates based on 16S rDNA sequence analysis.

| Phyla | Isolate | Closest Species Match | Identity (%) | Accession no. |
| --- | --- | --- | --- | --- |
| **<u>Adams Flat Isolate</u>** |  |  |  |  |
| Actinobacteria | VN1 | *Arthrobacter flavus* | 100 | MF541537.1 |
|  | PN6 | *Arthrobacter tumbae* | 99.8 | MK696231.1 |
|  | PN2 | *Arthrobacter tumbae* | 99.9 | KY753209.1 |
|  | N2 | *Streptomyces anulatus* | 100 | MK530154.1 |
|  | R8 | *Streptomyces badius* | 99.3 | MH482909.1 |
|  | S10 | *Streptomyces flavogriseus* | 99.8 | KX358684.1 |
|  | N3 | *Streptomyces pratensis* | 100 | MK484235.1 |
|  | N1 | *Streptomyces globisporus* | 100 | MF980904.1 |
|  | VN7 | *Streptomyces xinghaiensis* | 98.8 | KC493988.1 |
| Bacteroidetes | **VN2** | ***Belliella marina*** | **92.1** | **NR145847.1** |
|  | **VN3** | ***Cesiribacter roseus*** | **96.1** | **NR109022.1** |
|  | S14 | *Hymenobacter roseosalivarius* | 98.8 | NR029359.1 |
| Firmicutes | PN5 | *Bacillus luteolus* | 98.7 | KY849443.1 |
|  | PS1 | *Jeotgalibacillus marinus* | 99.6 | LR218079.1 |
|  | PN4 | *Jeotgalibacillus marinus* | 99.6 | LR218079.1 |
|  | VN4 | *Paenisporosarcina antarctica* | 99.7 | CP038015.1 |
|  | PN9 | *Paenisporosarcina quisquiliarum* | 98.7 | MH392666.1 |
|  | PN1 | *Paenisporosarcina quisquiliarum* | 99.5 | NR_043720.1 |
|  | VN5 | *Solicibacillus silvestris* | 100 | MH973485.1 |
| Proteobacteria | VR2 | *Altererythrobacter confluentis* | 99.3 | NR152077.1 |
|  | **PR3** | ***Roseomonas aquatica*** | **97.8** | **NR042501.1** |
| Total | 21 |  |  |  |
| **<u>Windmill Islands Isolate<sup>a</sup></u>** |  |  |  |  |
| Actinobacteria | NBH87 | *Frigoribacterium endophyticum* | 98.9 | NR_134732.1 |
|  | **NBWT11** | ***Klenkia marina*** | **97.2** | **NR_156810.1** |
|  | NBSH8 | *Pseudarthrobacter phenanthrenivorans* | 98.6 | NR_074770.2 |
|  | NBSH44 | *Streptomyces finlayi* | 99.9 | NR_043354.1 |
| Proteobacteria | **INR13** | ***Azosprillum zeae*** | **98** | **NR_043934.1** |
|  | **NBM11** | ***Novosphingobium stygium*** | **98.1** | **NR_040826.1** |
|  | NBH48 | *Paracoccus carotinifaciens* | 99.7 | NR_024658.1 |
|  | NBWT7 | *Sphingomonas jeddahensis* | 98.9 | NR_158139.1 |
| Total | 8 |  |  |  |

<sup>a</sup>Windmill Islands isolates were previously recovered and had their full genome 16S rDNA sequenced (N. Benaud, Unpublished data).
<sup>b</sup>Potential novel bacterial species recovered are in bold.

**Table 2.** Response of isolates to biosensors and HSL profiling of culture extracts.

| Phyla | Bacterial strain | NTL4<sup>a</sup> | CV026 | MT102 | Potential HSLs produced<sup>b</sup> |
| --- | --- | --- | --- | --- | --- |
| **<u>Adams Flat Isolate</u>** |  |  |  |  |  |
| Actinobacteria | *Arthrobacter sp.* VN1 | + | - | + | C6- , C9- , C11- |
|  | *Arthrobacter sp.* PN6 | - | - | + | C9- |
|  | *Arthrobacter sp.* PN2 | + | - | + | C8- |
|  | *Streptomyces sp.* N4 | - | - | - | - |
|  | *Streptomyces sp.* R8 | - | - | + | 3-oxo-C6- , C7- |
|  | *Streptomyces sp.* S10 | - | - | + | 3-oxo-C8- |
|  | *Streptomyces sp.* N3 | - | - | - | - |
|  | *Streptomyces sp.* N1 | + | - | + | 3-oxo-C6- , C7- |
|  | *Streptomyces sp.* VN7 | + | - | + | C7- , C8- , C10- |
| Bacteroidetes | *Belliella sp.* VN2 | + | - | - | N/A |
|  | *Cesiribacter sp.* VN3 | N/A | N/A | N/A | N/A |
|  | *Hymenobacter sp.* S14 | N/A | N/A | N/A | N/A |
| Firmicutes | *Bacillus sp.* PN5 | + | - | + | C7- , C8- , C10- |
|  | *Jeotgalibacillus sp.* PS1 | - | - | + | C8- |
|  | *Jeotgalibacillus sp.* PN4 | + | - | + | 3-oxo-C8- |
|  | *Paenisporosarcina sp.* VN4 | + | - | + | C7- |
|  | *Paenisporosarcina sp.* PN9 | - | - | - | - |
|  | *Paenisporosarcina sp.* PN1 | - | - | - | - |
|  | *Solicibacillus sp.* VN5 | + | - | + | C7- , C8- |
| Proteobacteria | *Altererythrobacter sp.* VR2 | N/A | N/A | N/A | N/A |
|  | *Roseomonas sp.* PR3 | N/A | N/A | N/A | N/A |
| Total |  | 9 | 0 | 12 |  |
| **<u>Windmill Islands Isolate</u>** |  |  |  |  |  |
| Actinobacteria | *Frigoribacterium sp.* NBH87 | - | - | + | C6- , C9- , C10- |
|  | *Klenkia sp.* NBWT11 | - | - | + | C6- |
|  | *Pseudarthrobacter sp.* NBSH8 | - | - | - | - |
|  | *Streptomyces sp.* SH44 | + | - | + | 3-oxo-C8- |
| Proteobacteria | *Azospirillum sp.* INR13 | + | - | + | C6- |
|  | *Novosphingobium sp.* NBM11 | - | - | + | C6- , C8- , 3-oxo-C10- , 3-oxo-C12- |
|  | *Paracoccus sp.* NBH48 | + | - | + | 3-oxo-C6- , C9, C10, C11, 3-oxo-C14- |
|  | *Sphingomonas sp.* NBWT7 | - | - | + | 3-oxo-C10- |
| Total |  | 3 | 0 | 7 |  |

N/A Isolates did not have significant growth to be included in this bioreporter assay
<sup>a</sup>Preliminary screening via plate bioassays using strains NTL4 & CV026; Small-batch ethyl acetate extraction using strain MT102
<sup>b</sup>Profiling of potential HSLs derived from large-batch ethyl acetate extraction, using strain MT102 as an agar overlay on TLC plates

Actinobacteria were dominant here (Figure 1), which is in line with knowledge of their inherent flexibility by virtue of harbouring wide metabolic capacities and survival mechanisms (12, 15, 20), thus giving them the selective advantage to dominate over other phyla in harsh environments. This phenomenon is not exclusive to just arid deserts but many other extreme environmental settings such as volcanic areas, dark caves and marine environments (65–67). The Actinobacteria community in these extreme habitats were found to produce various broad-spectrum antibiotics, enzymes, anti-tumor agents and immunomodulators. In fact, since discovery of the first antibiotic till today, 80% of the important drugs are sourced from members of Actinobacteria, mainly from the genus *Streptomyces* (68).

The majority recovered isolates exhibited 99-100% sequence similarity to characterised species (Table 1). Interestingly, potentially novel isolates were also recovered, including two Bacteroidetes members, *Belliella sp.* (92% similarity), *Cesiribacter sp.* (96%) and a Proteobacterium *Roseomonas sp.* (98%). The closest species match of isolates VN2, VN3 and PR3 were *Belliella marina, Cesiribacter roseus* and *Roseomonas aquatica,* all of which are gram-negative bacteria isolated from seawater, desert sand and drinking water, respectively (69–71). The low sequence similarity (92%) of isolate *Belliella sp.* VN2 suggests that it may represent a novel genus under the phylum Bacteroidetes. Further DNA-DNA whole genome hybridisation, fatty acid analysis along with a range of biochemical tests are required to confirm the novelty of these three new isolates.

Notable was the similarity of Soil 2 and Soil 3 in terms of the recovered taxa, whereas Soil 1 resulted in a distinct group of cultured members, primarily comprised of *Streptomyces* spp. (Figure 1). Soil environmental factors are known to effect microbial community structure and in fact, Soil 2 and Soil 3 had a similar alkaline pH of 8.7-8.8 (Supplementary Table 2) whereas Soil 1 had a pH of 7.8.

The two soil samples (Soil 2 and 3) with higher soil fertility, defined as organic carbon, nitrogen, and chloride content, had higher species richness over Soil 1. This means that the abundance of a certain bacterial species recovered in Soil 2 and Soil 3, were substantially higher than Soil 1 cultures, in which only two genera were successfully cultured along with low colony counts. The limited microbial growth observed for Soil 1 cultures could be associated with the markedly low phosphorus level (Supplementary Table 1), given that phosphorus is one of the vital limiting nutrients in polar soils (9, 72). These results were consistent with previous work on Antarctic soil biodiversity which used structural equation modelling to reveal that pH drives microbial community structure in polar soil ecosystems, that is, which taxa would be present, while soil fertility is positively correlated with bacterial species richness (9).

### HSL-based QS as a Universal Language within Antarctic Soil Microbes

Out of all 21 strain representatives recovered from Adams Flat, only 17 isolates were tested for HSL activity due to lack of significant growth in four isolates (*Hymenobacter sp.* S14*, Altererythrobacter sp.* VR2*, Cesiribacter sp.* VN3 and *Roseomonas sp.* PR3) under the pre-set laboratory conditions for this study. Along with the addition of eight Windmill Islands-recovered isolates, a total of 25 representative species were subjected to bioassays to determine HSL activity.

A diversity of presumptive HSL molecules was detected in 20/25 (80%) of grown to stationary phase, a stage with high cell densities, that is associated with nutrient depletion and accumulation of waste products that become toxic to the bacterial cells (73). Production of these signalling molecules under unfavourable conditions suggest that they play a role in regulating survival phenotypes. This explains why, production of secondary metabolites, biofilm formation and virulence, usually occurs during stationary phase.

For Adams flat-recovered isolates, 13/17 (76%) of the tested bacterial isolates were positive against at least one of the LuxI/LuxR-based QS biosensors. Amongst the eight Windmill Islands-recovered isolates, seven potential HSL producers were identified, three belonging to Actinobacteria, with four others Proteobacteria.

Surprisingly, within all bacterial isolates displaying active compound production, 15 were gram-positive bacteria spanning Actinobacteria and Firmicutes (Table 1). While HSL-based QS is common in gram-negative Proteobacteria (74), to our best knowledge, there are only four known reports of this QS system in gram-positive bacteria, two of which were of marine origin (36, 37), one was from mangrove rhizospheres (38) and one isolated from microbial mats (39). All the potential HSL producers isolated from these studies were also affiliated to either the Actinobacteria or Firmicutes phylum. The widespread discovery of potential HSL-based QS from bacteria spanning both gram-negative and gram-positive in Antarctic soils suggests HSL-based QS systems may be a universal language for communication by soil microbes.

It was previously thought that QS in gram-positive bacteria relies on short peptides instead of HSLs as signalling molecules (75, 76) but, subsequently genes encoding for peptide signals were found in gram-negative bacteria via *in silico* analyses (77) and HSL activity has been reported in an expanding breadth of the gram-positive bacterial community (36–39). With different species colonising the same habitat in proximity, there is no doubt QS regulatory cassette can be transmitted between organisms via horizontal gene transfer. This phenomenon suggested that bacteria may have co-evolved to be multi-lingual with the presence of multiple QS systems, so that communication at multiple distinct levels: intra-species, inter-species within the same genus, inter-species between different genera and even inter-domain, is made easier (33–35).

### Profiling of presumptive HSL via TLC

The extracted active compounds separated on thin layer chromatogram were identified according to *Rf* values calculated based on distance migrated by the molecules, along with tailing properties in comparison to a range of synthetic HSL standards.

Several active compounds appear on the TLC sheet after 8 h of incubation at 30 °C while some compounds were only visible after 18 h. These presumptive HSLs became oversaturated at 18 hr, with several new compounds only becoming visible at this time point (Figure 2). However, this does not reflect the concentration of a certain type of HSL being produced as the biosensor *E. coli* MT102 has different sensitivities against different HSLs. For instance, 0.009 ng of OHHL can induce green fluorescence by the biosensor while 113.2 ng of C12-HSL is required to induce the same amount of fluorescence (48). None of the 3-oxo-derivatives, including those made synthetically, had tailing properties as proposed in previous literature (50), except for N-(3-oxodecanoyl)-L-HSL (ODHL).

**Figure 2.**
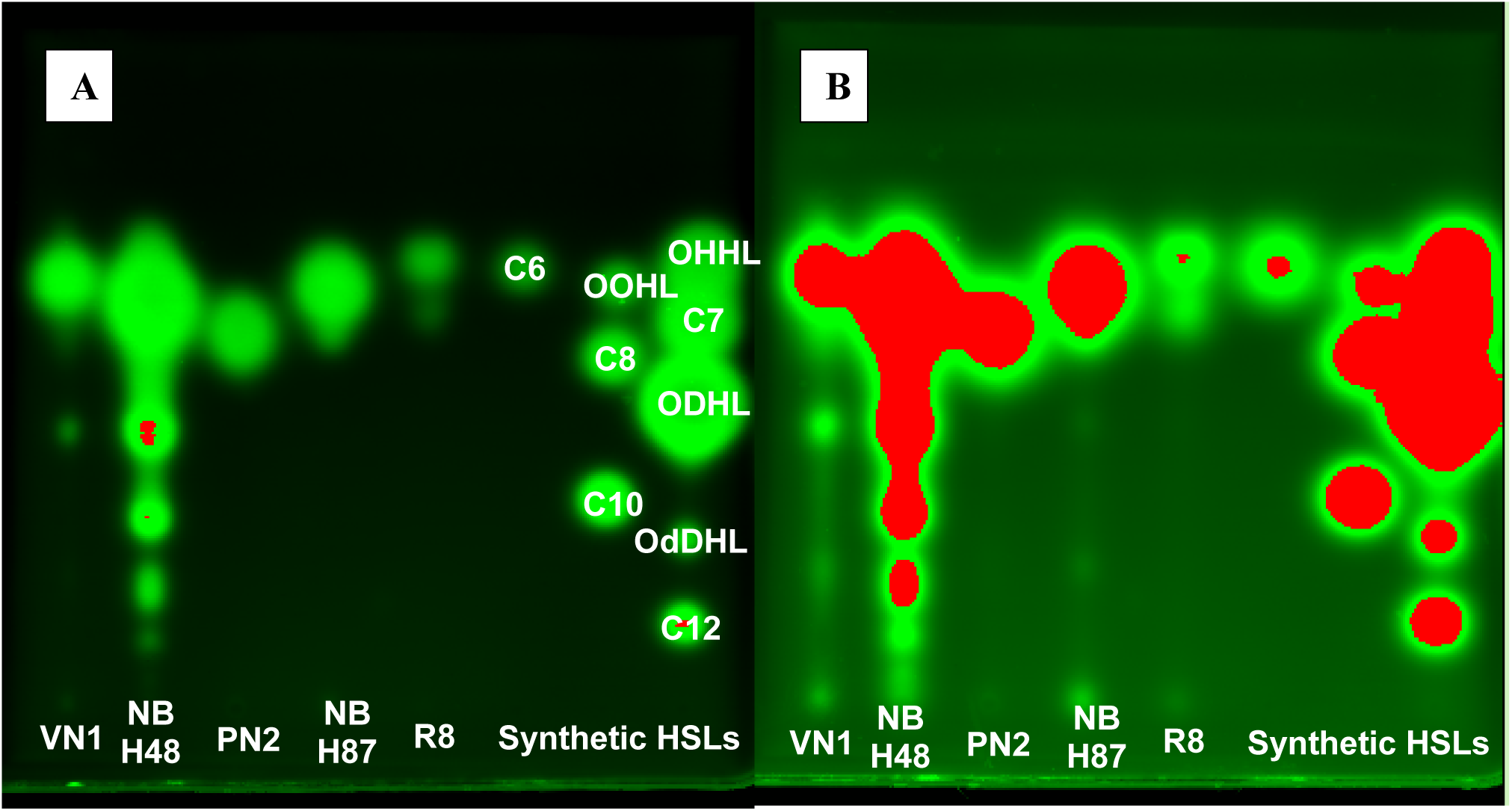
Visualisation of active compounds with agar overlay of MT102 bioreporter incubated at 30 °C for (A) 8 h (B) 18 h. Presumptive HSLs extracted from cell-free culture supernatants were separated by TLC developed with methanol/water (60:40 *v/v*). Culture extracts for isolates VN1, NBH48, PN2, NBH87, R8, and positive controls are presented (left to right).

All detectable HSLs extracted in the present study were of medium to long chain length, with C6-being the shortest and C14-being the longest chain HSL extracted. A relatively long chain length may be required to resist HSL hydrolysis that has been observed previously in alkaline conditions (78, 79), especially given the alkaline pH of the soil in the present study (Supplementary Table 1). C7-HSL appeared to be the potential signalling molecule of highest occurrence and was produced by seven different isolates, affiliated to three phyla, namely, Actinobacteria, Firmicutes and Proteobacteria (Table 1). The active compound produced by a Bacteroidetes isolate *Belliella sp.* VN2 was detected by biosensor *A. tumefaciens* NTL4 but not by *E. coli* MT102, suggesting that the active compounds may be 3-hydroxy-derivatives of HSL which cannot be detected by *E. coli* MT102. However, it should be noted that confirmation of signal molecule identity would entail employing mass spectrometry and nuclear magnetic resonance in follow-up studies.

Active compounds detected in the cultures of gram-positive bacteria ranged from C6 to C11-HSLs, with no evidence of the short chain C4-HSL. Gram-positive bacteria that were identified as potential HSL-producers from previous studies (36–38), also produced medium to long chain length active compounds ranging from C8 to C14-HSLs, albeit the non-alkaline pH of their respective environmental setting, except for one Actinobacterium isolated from a hypersaline microbial mat, which was found to produce C4-HSL (39). These findings may indicate gram-positive bacteria, if capable of producing HSLs, are not in favour of short-chain HSLs, or these compounds could be produced in concentrations ineffective for standard bioassays.

Members of Proteobacteria yielded the highest number of detectable compounds, with the production of a minimum of four potential HSLs by *Paracoccus sp.* NBH48 and five potential HSLs by *Novosphingobium sp.* NBM11. This is not surprising given that LuxI/LuxR-based QS is common within Proteobacteria, and hence is able to make full use of this communication system. The highest production of presumptive HSLs in Proteobacterial isolates was also reported in another study (38), compared to members of Actinobacteria and Firmicutes, which are relatively unfamiliar with this language. Despite the relatively high number of potential HSLs produced by *Novosphingobium sp.* NBM11, they were not detectable with the broad range-detectivity biosensor *A. Tumefaciens* NTL4.

### Genome sequences of Windmill Islands isolates

Of the eight genomes examined, six were high-quality assemblies, displaying high contiguity (L50=1; N50>3 Mb), and completeness (>97% complete) (Supplementary Table 1). They were the *Streptomyces sp.* NBSH44, *Sphingomonas sp.* NBWT7, *Pseudoarthrobacter sp.* NBSH8, *Klenkia sp..* NBWT11, *Novosphingobium sp.* NBM11, and *Frigoribacterium sp.* NBH87. The genomes of *Paracoccus sp.* NBH48 and *Azosprillum sp.* INR13 diverged from high-quality indicator range for completeness (83% and 65% completion, respectively), and exhibited fragmentation (12 and 50 contigs, respectively) and low coverage (22 and 18, respectively).

### Genetic basis for QS in Antarctic isolates

Despite active detection of signalling molecules in bioreporter assays in the present study, LuxI/LuxR homologs were not found in genomes of all these isolates. By searching for HSL synthase COG category COG3916 and the IPR codes of domains associated with autoinducer synthase (Supplementary Table 3), we identified one LuxI homolog each in the gram-negative Proteobacteria genomes of *Paracoccus sp.* NBH48 (A1PzeaV1_2917) and *Novosphingobium sp.* NBM11 (A2NstyV1_00719). No *luxI* genes were found in any of the available genomes of gram-positive Actinobacteria, despite active HSL production detected in the *E.coli* MT102 bioassay. The same scenario has been reported by (39), where no HSL synthase genes were identified within gram-positive genomes of Actinobacteria and Firmicutes. The remaining two Proteobacterial isolates (NBR13 and NBWT7) that showed HSL activity in the bioreporter assays, had no signs of *luxI* genes in their genomes, suggesting that these LuxI/LuxR sequences may not be very well conserved in Antarctic isolates due to adaptation and evolution under constant environmental stressors.

Based on PROKKA-annotation, the LuxI homolog found in isolate NBM11 (A2NstyV1_00719) is an isovaleryl-HSL synthase, which synthesises an unusual branched-chain HSL signal (29). Phylogenetic analyses (Figure 3) further suggest that the *Novosphingobium sp.* M11 LuxI homolog may represent a novel autoinducer synthase that does not produce the common acyl-HSL, as shown by it clustering with the CoA-utilizing autoinducer synthases that encode for atypical HSL signals. This may explain the lack of detection of HSL production by the biosensors CV026 and NTL4, which only detect normal acyl-HSLs (28, 29, 80). However, this does not exclude the possibility that HSL signals were produced in levels beyond biosensor NTL4 detection limits and thus resulted in false negative. On the other hand, LuxI sequence retrieved from isolate *Paraococcus sp.* NBH48 (A1PzeaV1_2917) clustered with that of *Paracoccus zhejiangensis,* suggesting conservation within the same genus.

**Figure 3.**
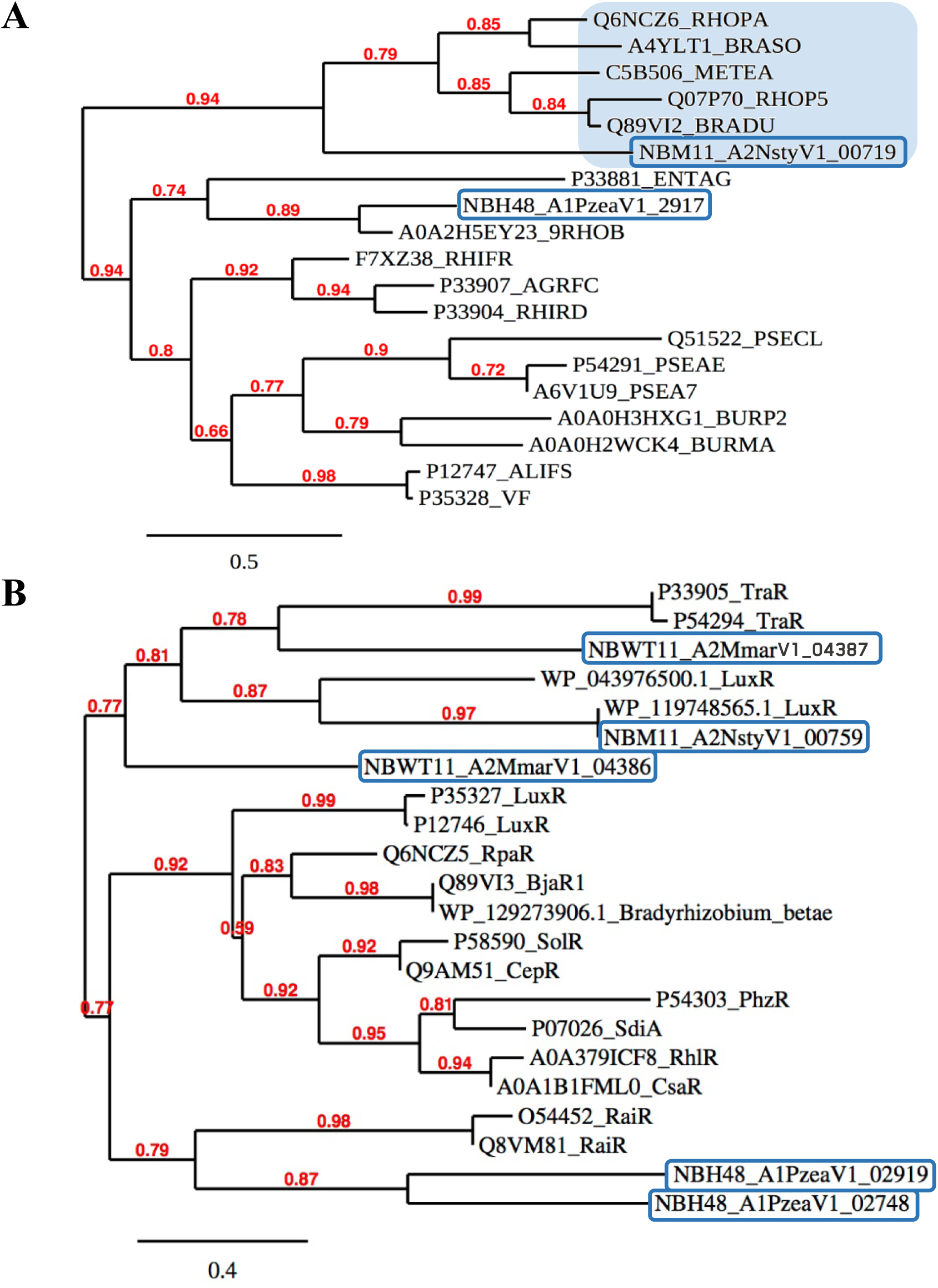
Protein phylogeny of (A) HSL synthases (B) LuxR regulators. The clade shaded in blue are HSL synthases that produce atypical HSL signals from coenzyme-A (CoA)-activated acids instead of the usual acylated-acyl carrier protein-activated acid. LuxI and LuxR homologs retrieved from Windmill Islands isolate genomes are within blue boxes. These maximum-likelihood trees were constructed with reference amino acid sequences of HSL synthases and LuxR homologs retrieved from UniProtKB and NCBI database. Numbers at nodes indicate bootstrap probabilities of 1000 bootstrap re-samplings. Bar, 0.5 and 0.4 substitutions per amino acid position for Phylogenetic tree (A) and (B), respectively.

Multiple sequence alignments of putative *luxI* in *Paraococcus sp.* NBH48 (A1PzeaV1_2917) and *Novosphingobium sp.* NBM11 (A2NstyV1_00719) against *Vibrio fischeri* resulted in low identity similarity, 24.53% and 38.27%, respectively (Supplementary Figure 2). However, the eleven critical residues reported to be key sites in LuxI family proteins (81): R-25, E-44, D-46, D-49, D-60, G-67, R-70, R-104, A-133, E-150 and G-164, were conserved in these two LuxI sequences, except for D-46, D-60, A-133 and E-150 in isolate NBH48. LuxI sequence found in isolate NBM11 showed conservation in all the eleven key residues except for E-150, which was reported to be less well-conserved (81). Thus, these LuxI homologs are still able to carry out critical enzymatic activity required to catalyse HSL production regardless the low conservation consensus and using a different substrate, coenzyme-A (CoA)-activated acid, instead of the usual acylated-acyl carrier protein-activated acid to synthesise HSLs.

A total of 135 potential LuxR sequences were extracted by searching for LuxR-associated IPR codes (Supplementary Table 3). However, only five of these, derived from three genomes, were predicted to have the ability to bind HSL ligands as determined by COACH. As such, these five sequences were the only LuxR sequences subjected to further analysis, in order to exclude other non-LuxR sequences that were picked up during IPR code search due to the common HTH domain they possess. LuxR homologs are problematic to retrieve from full genomes due to the generic domain it possesses, a DNA-binding, helix-turn-helix (HTH) domain which is also present in numerous non-LuxR regulators (82). Horizontal gene transfer between microorganisms could further complicate genome mining as this could result in increased variabilities (and thus decreased homology) in the LuxI/LuxR homologs due to permutation and combination.

Within the three genomes in which LuxR homologs were found, three LuxR homologs were in the gram-negative Proteobacteria isolates, *Paracoccus sp.* NBH48 (A1PzeaV1_2919 and A1PzeaV1_02748) and *Novosphingobium sp.* NBM11 (A2NstyV1_00759), while two LuxR homologs were derived from the gram-positive Actinobacteria member, *Klenkia sp.* NBWT11 (A2MmarV1_04386 and A2MmarV1_04387).

Multiple sequence alignments of these putative LuxR homologs against *V. fischeri* displayed conservation in a number of amino acid residues, including those ones that were identified as important residues in LuxR regulatory activity (83), such as W-66, Y-70, D-79, P-80, V-109, G-117, G-121, L-183, T-184, R-186, E-187, L-191, A-192, G-197, I-203, I-206, L-207, T-213, V-214, H-217, L-218, K-224 and R-230 (Supplementary Figure 3). Conservation of these critical residues in *luxR* sequences of the gram-positive isolate *Klenkia sp.* WT11 showed that LuxR homologs of gram-positive bacteria are similar to that of gram-negative bacteria, further emphasising that horizontal gene transfer is the likely source of acquisition of this QS cassette in gram-positive community.

### Potential traits regulated by LuxI/LuxR

Genes in the vicinity of putative LuxI/LuxR homologs were found to encode for functions that provide survival advantage.The *luxI/luxR* cassette in NBH48 (A1PzeaV1_2917/A1PzeaV1_2919) were found in separate strands, in a convergent direction I⃗R⃖(Table 3), with virulence and heavy metal transporter genes downstream of each *lux* genes in their respective contigs. The virulence factor virB4 that was identified here, has been shown to increase cell survival by inhibiting apoptosis upon stimuli (84) and mediates plasmid conjugation (85). Mercury and copper transporters are known to confer resistance to heavy metals that can be toxic to the bacteria. There was no LuxI homolog found near the other LuxR homolog in NBH48 (A1PzeaV1_02748) but a cardiolipin synthetase was adjacent to it, on the opposite strand. Cardiolipin has been reported to mediate various signalling pathways (86), and thus, may be working in tandem with this LuxR homolog in I⃖R⃖ arrangement.

**Table 3.**
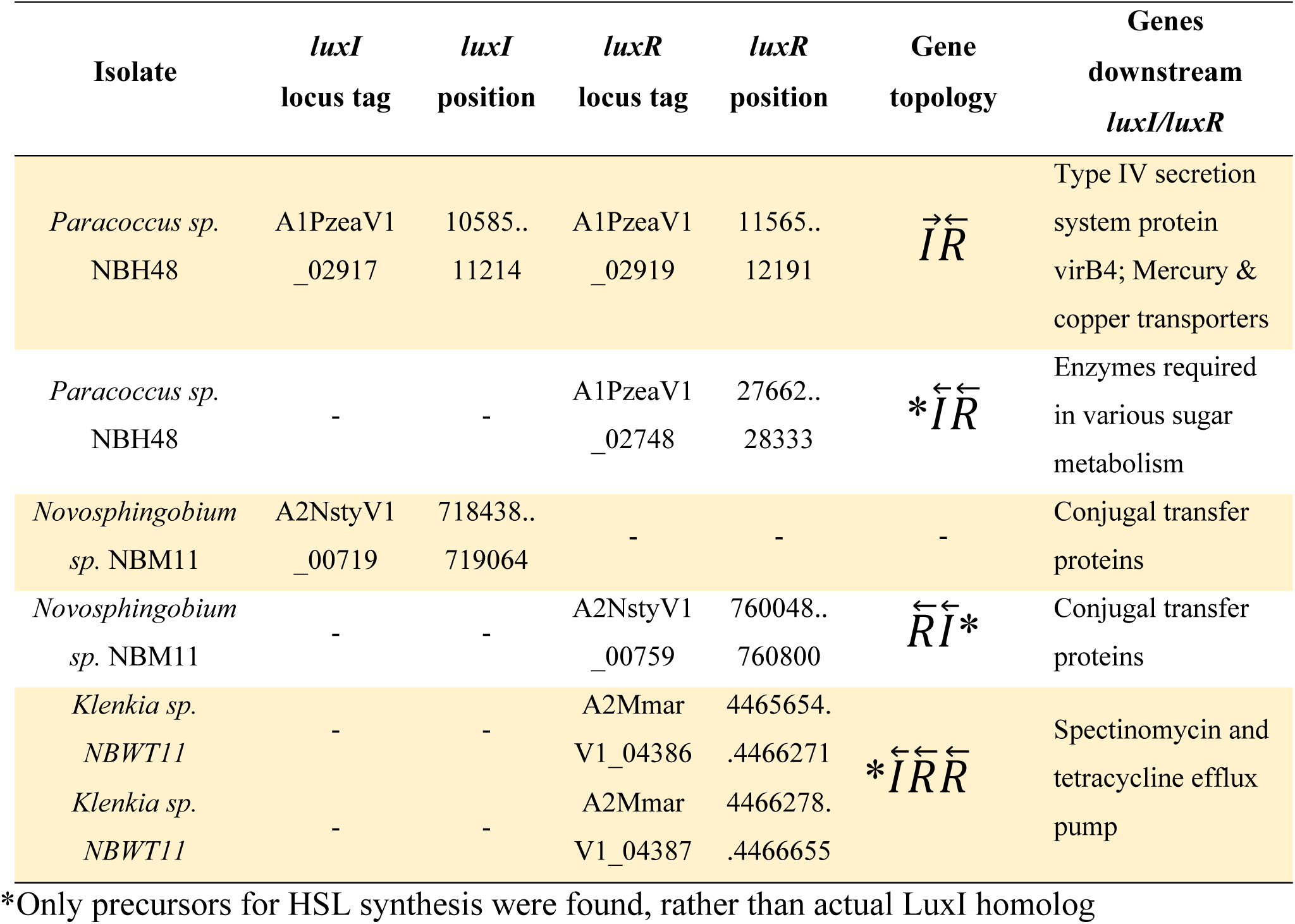
Topology of *luxI/luxR* genes in Windmill Island isolates.

The putative LuxI and LuxR homologs found in NBM11 were 42 000 nt apart, suggesting they may not be working in tandem. The putative *luxI* in NBM11 (A2NstyV1_00719) were flanked by several genes encoding for conjugal transfer proteins, which are essential for passing on useful traits between microorganisms. No *luxR* gene was observed in vicinity to the NBM11 *luxI,* but a HTH-type domain transcriptional regulator, MarR, was found upstream. As for the NBM11 putative *luxR* (A2NstyV1_00759), a set of precursors of HSL synthesis such as long chain fatty acid-CoA ligase and a few different acyl-CoA substrates, were found upstream, in a R⃖I⃖ topology. Downstream were proteins involved in conjugation, suggesting NBM11 *lux* genes were located on a conjugal plasmid, similar to *traI/traR* in *A. tumefaciens* (87).

In NBWT11, two LuxR homologs (A2MmarV1_04386 and A2MmarV1_04387) were found on the antisense strand in R⃖R⃖ topology. There was no LuxI homolog but an acetyl-CoA synthetase and a spectinomycin and tetracycline transporter were located downstream of the double LuxR homologs. This multiantibiotic efflux pump can confer antimicrobial resistance, enhancing the bacteria competitive advantage (88). However, further experimental work is required to determine if these genes were actively regulated by the adjacent *luxI/luxR* genes.

## CONCLUSION

Bacteria surviving in East Antarctic soils are exposed to a number of environmental threats that must drive a variety of survival responses that require cell-to-cell communication. This study reports for the first time, quorum sensing activity within Antarctic soils with profiles of the active signalling molecules produced by both gram-negative and gram-positive bacteria, the latter which has only been recently discovered to be capable of this new communication system. As Antarctic microbes represent a reservoir for novel functionalities and natural products, understanding the basis of microbial communication, which is a pre-requisite to accomplish almost any processes, is one of the first steps to manipulate microbes to our benefits.

This study has provided a platform for future studies further characterising the physiological adaptations of microorganisms that allow them to not just survive but thrive under extreme Antarctic conditions.

## ACKNOWLEDGEMENTS

We thank the Australian Antarctic Program and expedition teams in 2012 for sampling of Antarctic soils used in this study. We thank Bioplatforms Australia for the provision of Vestfold Hills biodiversity data [https://data.bioplatforms.com/organization/about/australian-microbiome]. This work was supported by the University International Postgraduate Award (UIPA) (awarded to S.Y.W), the Australian Research Council Future Fellowship (FT170100341; awarded to B.C.F). Special thanks to Dr. Onder Kimyon for providing us the bioreporter strains and Eden Zhang for the bubble plot.

## SUPPLEMENTARY MATERIALS

### Supplementary Figures

**Supplementary Figure 1.**
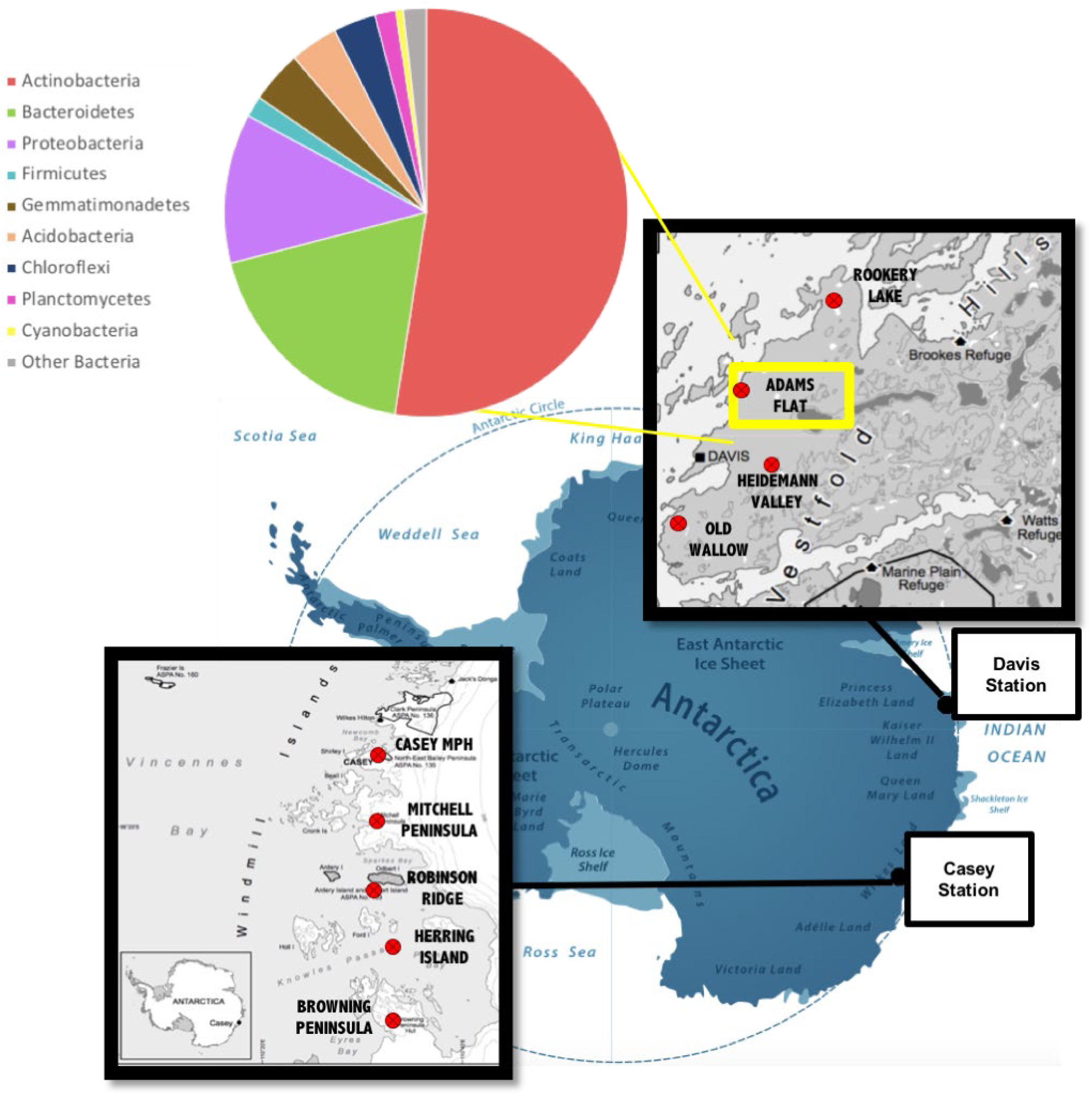
Geographical map of Windmill Islands (left) and Vestfold Hills (right), along with Adams Flat soil microbial community structure. Adams Flat from Vestfold Hills region was selected for cultivation-dependent analysis in this study. Community structure was determined via cultivation-independent 16S rDNA sequencing on Adams Flat soil samples that showed active trace gas chemosynthesis (41).

**Supplementary Figure 2.**
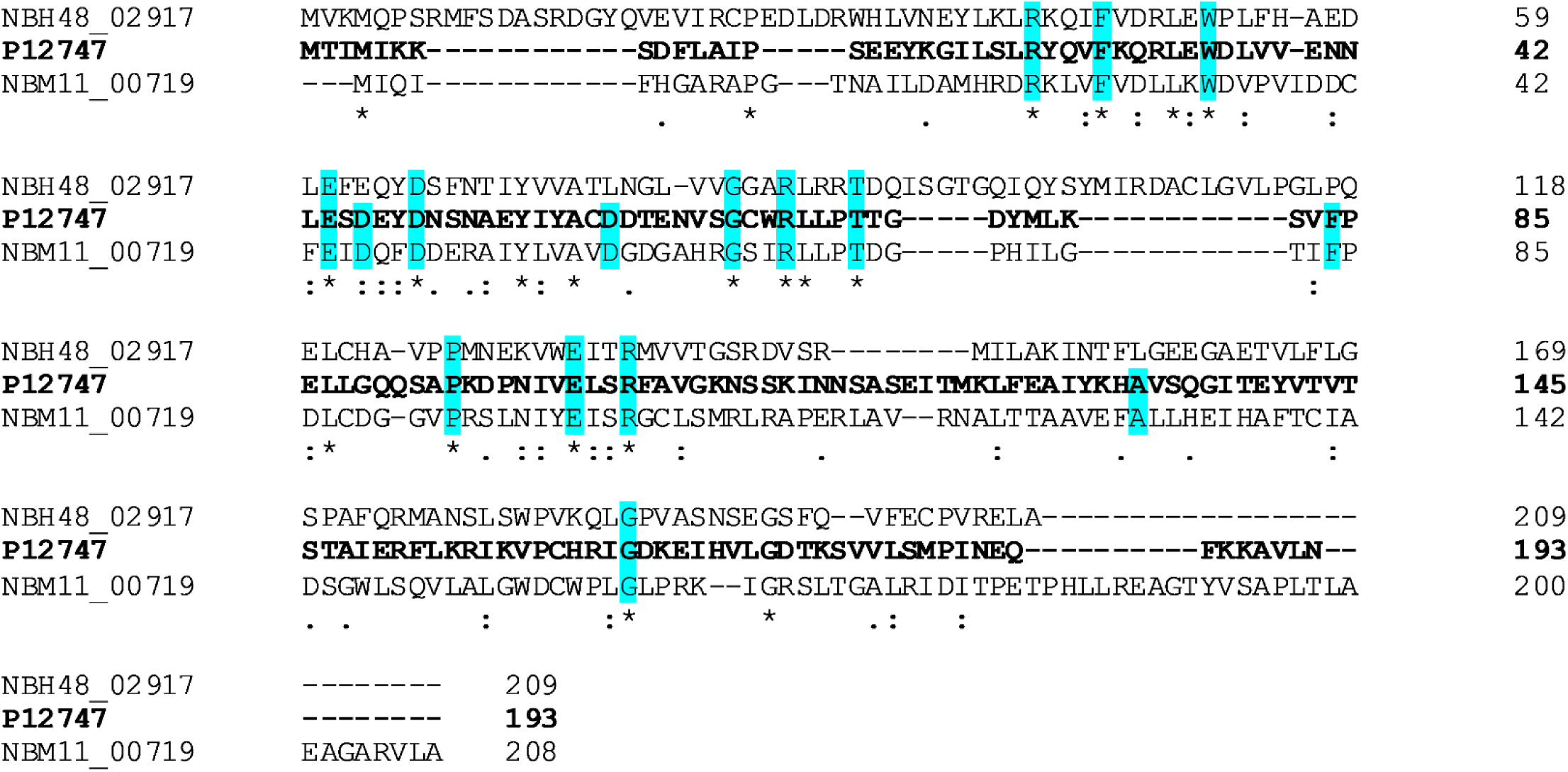
Multiple sequence alignment of LuxI sequences of *Paracoccus sp.* NBH48 and *Novosphingobium sp.* NBM11 against *V*. *fischerii* LuxI via Clustal Omega. Highlighted in blue are critical amino acid residues for LuxI activity (81, 89). * (asterisk) indicates fully conserved residue. : (colon) indicates conservation between groups of strongly similar amino acids – scoring > 0.5 in the Gonnet PAM 250 matrix. . (period) indicates conservation between groups of weakly similar properties – scoring =< 0.5 in the Gonnet PAM 250 matrix

**Supplementary Figure 3.**
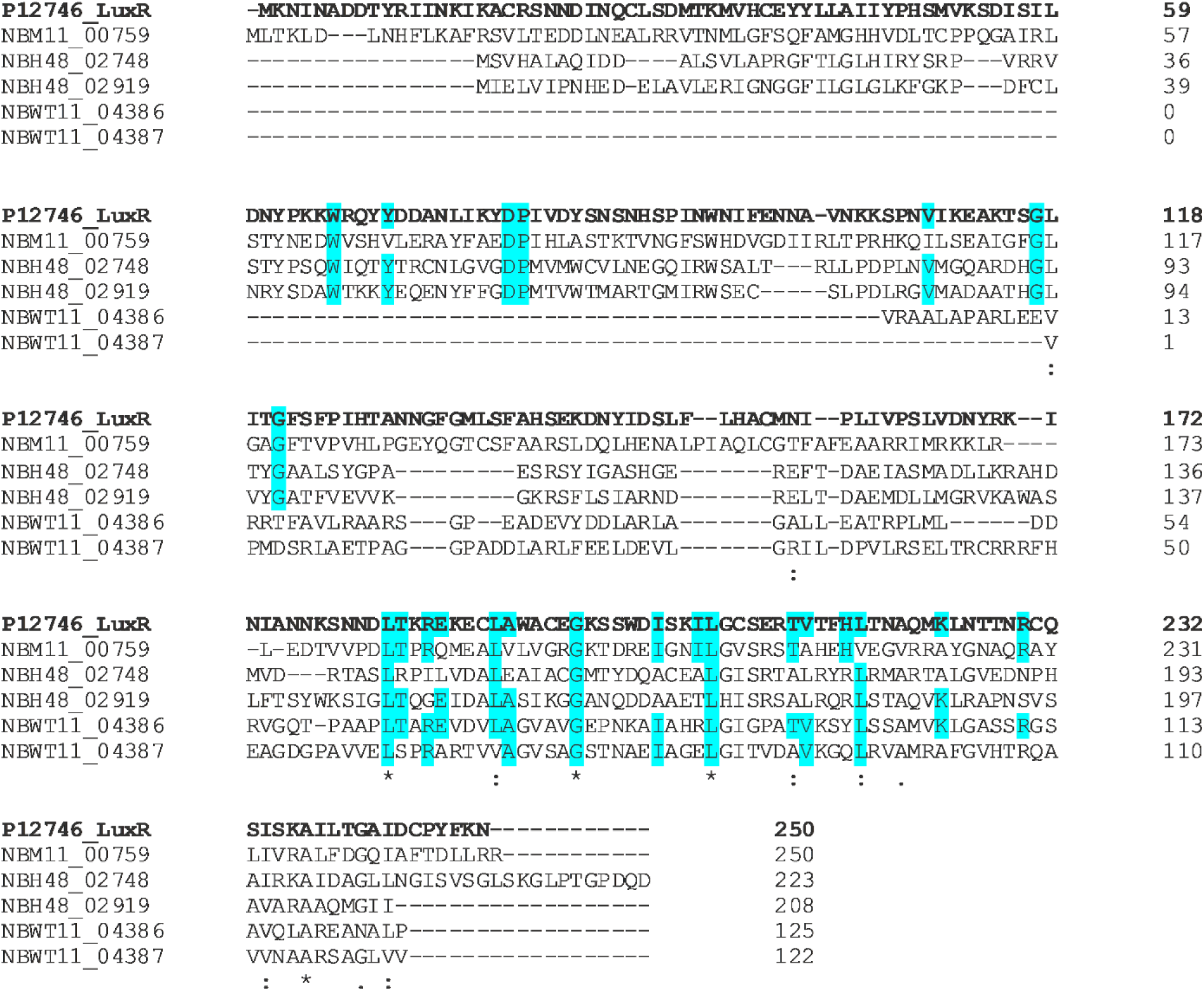
Multiple sequence alignment of LuxR sequences of *Paracoccus sp.* NBH48, *Novosphingobium sp.* NBM11 and *Klenkia sp.* NBWT11 against *V*. *fischerii* LuxR via Clustal Omega. Highlighted in blue are critical residues for LuxR activity (83, 89). * (asterisk) indicates fully conserved residue. : (colon) indicates conservation between groups of strongly similar amino acids – scoring > 0.5 in the Gonnet PAM 250 matrix . (period) indicates conservation between groups of weakly similar properties – scoring =< 0.5 in the Gonnet PAM 250 matrix

### Supplementary Tables

**Supplementary Table 1.**

| Strain | Assembled size (Mb) | Contigs | L50 | N50 (Mb) | Mean Coverage | Complete (%) | Contam (%) | CDS | tRNA | tmRNA | rDNA | No. of CDS assigned to COG/ (% of all CDS) |
| --- | --- | --- | --- | --- | --- | --- | --- | --- | --- | --- | --- | --- |
| *Streptomyces* NBSH44 | 7.7 | 3 | 1 | 7.47 | 94 | 99.7 | 0.7 | 7021 | 82 | 1 | 18 | 4962 (70.6) |
| *Paracoccus* NBH48 | 3.0 | **12** | **2** | 0.71 | **22** | **82.6** | 0.4 | 3053 | 47 | 0 | 9 | 2576 (84.4) |
| *Klenkia* NBWT11 | 4.6 | 1 | 1 | 4.59 | 33 | 97.7 | 0.8 | 4441 | 53 | 1 | 9 | 3595 (81.0) |
| *Novosphingobium* NBM11 | 5.3 | 4 | 1 | 4.15 | 26 | 99.1 | 1.9 | 4996 | 60 | 0 | 6 | 3909 (78.2) |
| *Sphingomonas* NBWT7 | 3.4 | 2 | 1 | 3.26 | 45 | 99.6 | 0.7 | 3231 | 53 | 0 | 6 | 2673 (82.7) |
| *Azospirillum* INR13 | 4.7 | **50** | **10** | **0.13** | **18** | **64.5** | 4.3 | 4447 | 65 | 0 | 24 | 3573 (80.3) |
| *Pseudarthrobacter* NBSH8 | 4.1 | 1 | 1 | 4.06 | 64 | 99.7 | 0.2 | 3745 | 53 | 1 | 12 | 3085 (82.4) |
| *Frigoribacterium* NBH87 | 3.4 | 2 | 1 | 3.33 | 37 | 98.5 | 0 | 3146 | 54 | 1 | 6 | 2431 (77.3) |

Numbers in bold indicate lower quality markers

**Supplementary Table 2.**
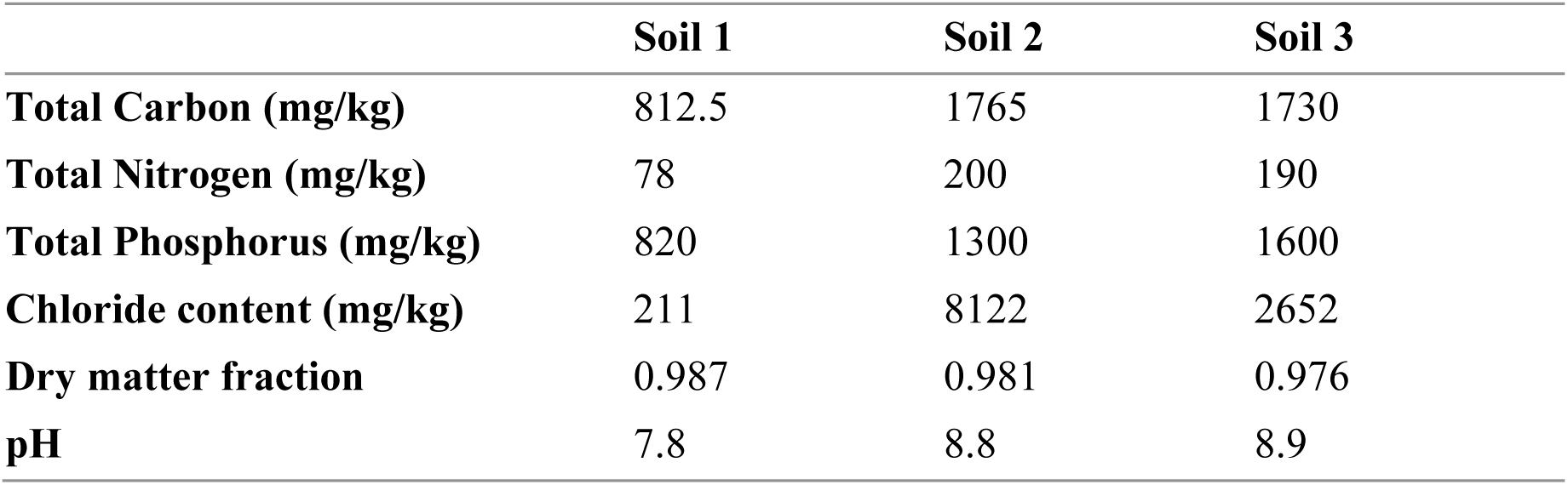
Measured soil parameters for the three selected soil samples.

**Supplementary Table 3.**
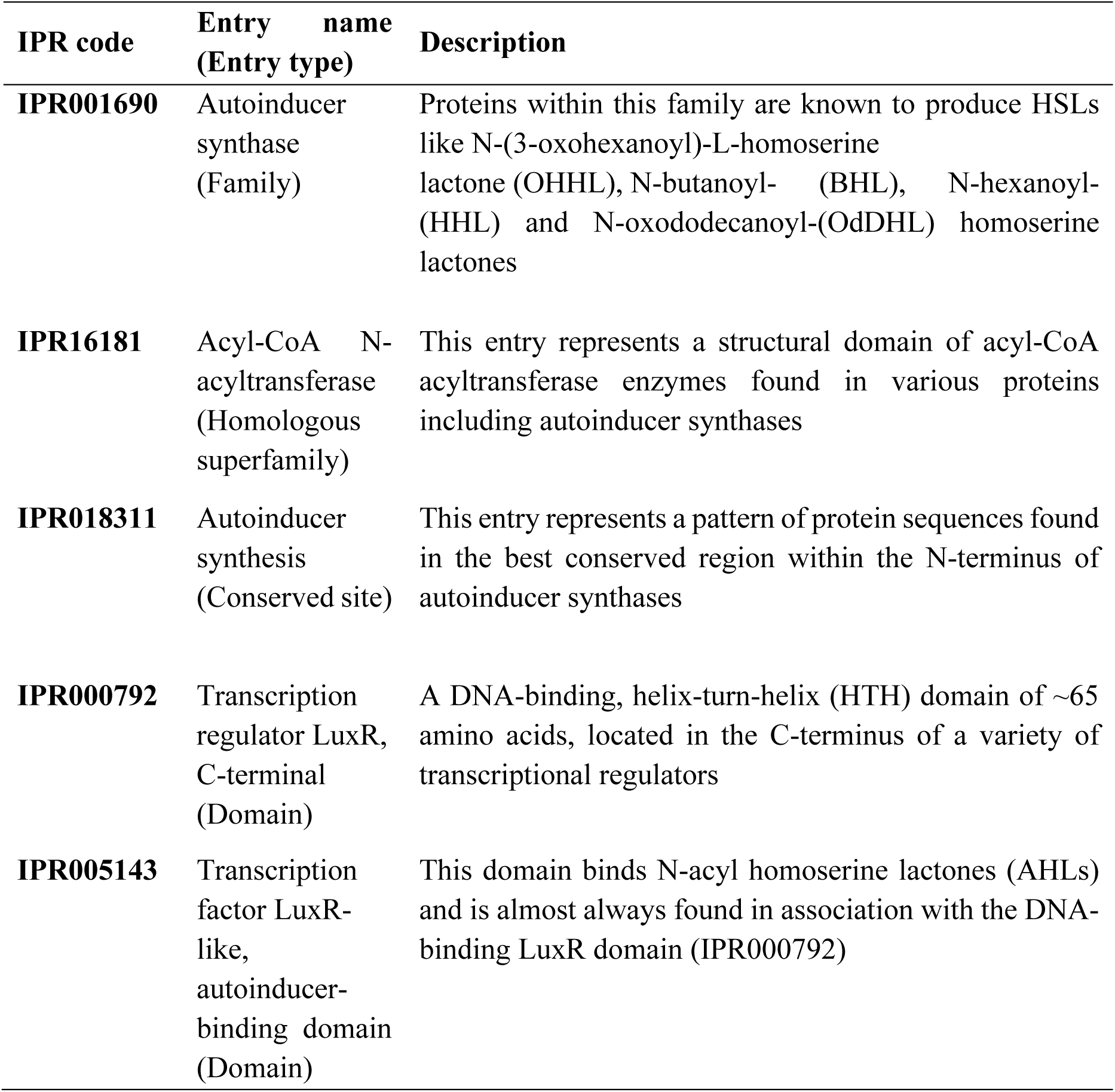
LuxI/LuxR-associated IPR codes searched against genome

| **IPR code** | **Entry name**<br>**(Entry type)** | **Description** |
| --- | --- | --- |
| **IPR001690** | Autoinducer synthase (Family) | Proteins within this family are known to produce HSLs like N-(3-oxohexanoyl)-L-homoserine lactone (OHHL), N-butanoyl- (BHL), N-hexanoyl- (HHL) and N-oxododecanoyl-(OdDHL) homoserine lactones |
| **IPR16181** | Acyl-CoA N-acyltransferase (Homologous superfamily) | This entry represents a structural domain of acyl-CoA acyltransferase enzymes found in various proteins including autoinducer synthases |
| **IPR018311** | Autoinducer synthesis (Conserved site) | This entry represents a pattern of protein sequences found in the best conserved region within the N-terminus of autoinducer synthases |
| **IPR000792** | Transcription regulator LuxR, C-terminal (Domain) | A DNA-binding, helix-turn-helix (HTH) domain of ~65 amino acids, located in the C-terminus of a variety of transcriptional regulators |
| **IPR005143** | Transcription factor LuxR-like, autoinducer-binding domain (Domain) | This domain binds N-acyl homoserine lactones (AHLs) and is almost always found in association with the DNA-binding LuxR domain (IPR000792) |

## REFERENCE

1. Fernandez Carazo R. 2011. Contribution to the study of past and present cyanobacterial diversity in Antarctica. PhD thesis. University of Liege, Belgium.

2. van Dorst J, Benaud N, Ferrari B. 2017. New Insights into the Microbial Diversity of Polar Desert Soils: A Biotechnological Perspective, p 169–183, Microbial Ecology of Extreme Environments doi:10.1007/978-3-319-51686-8_7.

3. Cullather RI, Bromwich DH, Woert MLV. 1998. Spatial and Temporal Variability of Antarctic Precipitation from Atmospheric Methods. J Clim 11:334–367.

4. Brandt RE, Warren SG. 1993. Solar-heating rates and temperature profiles in Antarctic snow and ice. J Glaciol 39:99–110.

5. Turner J, Anderson P, Lachlan-Cope T, Colwell S, Phillips T, Kirchgaessner A, Marshall GJ, King JC, Bracegirdle T, Vaughan DG, Lagun V, Orr A. 2009. Record low surface air temperature at Vostok station, Antarctica. 114.

6. Cornelius PE. Life in Antarctica, p 9–14. *In* (ed), Springer New York,

7. Cowan DA, Makhalanyane TP, Dennis PG, Hopkins DW. 2014. Microbial ecology and biogeochemistry of continental Antarctic soils. Front Microbiol 5:154.

8. Bottos EM, Scarrow JW, Archer SDJ, McDonald IR, Cary SC. 2014. Bacterial Community Structures of Antarctic Soils, p 9–33, Antarctic Terrestrial Microbiology doi:10.1007/978-3-642-45213-0_2.

9. Siciliano SD, Palmer AS, Winsley T, Lamb E, Bissett A, Brown MV, van Dorst J, Ji M, Ferrari BC, Grogan P, Chu H, Snape I. 2014. Soil fertility is associated with fungal and bacterial richness, whereas pH is associated with community composition in polar soil microbial communities. Soil Biol Biochem 78:10–20.

10. Cary SC, McDonald IR, Barrett JE, Cowan DA. 2010. On the rocks: the microbiology of Antarctic Dry Valley soils. Nat Rev Microbiol 8:129–38.

11. Ferrari BC, Bissett A, Snape I, van Dorst J, Palmer AS, Ji M, Siciliano SD, Stark JS, Winsley T, Brown MV. 2016. Geological connectivity drives microbial community structure and connectivity in polar, terrestrial ecosystems. Environ Microbiol 18:1834–49.

12. Ji M, van Dorst J, Bissett A, Brown MV, Palmer AS, Snape I, Siciliano SD, Ferrari BC. 2015. Microbial diversity at Mitchell Peninsula, Eastern Antarctica: a potential biodiversity “hotspot”. Polar Biol 39:237–249.

13. Gonzalez-Rocha G, Munoz-Cartes G, Canales-Aguirre CB, Lima CA, Dominguez-Yevenes M, Bello-Toledo H, Hernandez CE. 2017. Diversity structure of culturable bacteria isolated from the Fildes Peninsula (King George Island, Antarctica): A phylogenetic analysis perspective. PLoS One 12:e0179390.

14. Yergeau E, Bokhorst S, Huiskes AH, Boschker HT, Aerts R, Kowalchuk GA. 2007. Size and structure of bacterial, fungal and nematode communities along an Antarctic environmental gradient. FEMS Microbiol Ecol 59:436–51.

15. Benaud N, Zhang E, van Dorst J, Brown MV, Kalaitzis JA, Neilan BA, Ferrari BC. 2019. Harnessing long-read amplicon sequencing to uncover NRPS and Type I PKS gene sequence diversity in polar desert soils. FEMS Microbiol Ecol 95.

16. Dieser M, Greenwood M, Foreman CM. 2010. Carotenoid Pigmentation in Antarctic Heterotrophic Bacteria as a Strategy to Withstand Environmental Stresses. Arct Antarct Alp Res 42:396–405.

17. Hibbing ME, Fuqua C, Parsek MR, Peterson SB. 2010. Bacterial competition: surviving and thriving in the microbial jungle. Nat Rev Microbiol 8:15–25.

18. O’Brien J, Wright GD. 2011. An ecological perspective of microbial secondary metabolism. Curr Opin Biotechnol 22:552–8.

19. Chilton AM, Neilan BA, Eldridge DJ. 2017. Biocrust morphology is linked to marked differences in microbial community composition. Plant Soil 429:65–75.

20. Makhalanyane TP, Valverde A, Gunnigle E, Frossard A, Ramond JB, Cowan DA. 2015. Microbial ecology of hot desert edaphic systems. FEMS Microbiol Rev 39:203–21.

21. Williams L, Borchhardt N, Colesie C, Baum C, Komsic-Buchmann K, Rippin M, Becker B, Karsten U, Büdel B. 2016. Biological soil crusts of Arctic Svalbard and of Livingston Island, Antarctica. Polar Biol 40:399–411.

22. Davati N, Najafi MBH. 2013. Overproduction Strategies for Microbial Secondary Metabolites, p 23–37. In (ed), International Journal of Life Science & Pharma Research.

23. Rossi F, De Philippis R. 2015. Role of cyanobacterial exopolysaccharides in phototrophic biofilms and in complex microbial mats. Life (Basel) 5:1218–38.

24. Vásquez-Ponce F, Higuera-Llantén S, Pavlov MS, Ramírez-Orellana R, Marshall SH, Olivares-Pacheco J. 2017. Alginate overproduction and biofilm formation by psychrotolerant Pseudomonas mandelii depend on temperature in Antarctic marine sediments. Electron J Biotechnol 28:27–34.

25. Bihary D, Toth M, Kerenyi A, Venturi V, Pongor S. 2014. Modeling bacterial quorum sensing in open and closed environments: potential discrepancies between agar plate and culture flask experiments. J Mol Model 20:2248.

26. Decho AW, Norman RS, Visscher PT. 2010. Quorum sensing in natural environments: emerging views from microbial mats. Trends Microbiol 18:73–80.

27. Montgomery K, Charlesworth JC, LeBard R, Visscher PT, Burns BP. 2013. Quorum sensing in extreme environments. Life (Basel) 3:131–48.

28. Schaefer AL, Greenberg EP, Oliver CM, Oda Y, Huang JJ, Bittan-Banin G, Peres CM, Schmidt S, Juhaszova K, Sufrin JR, Harwood CS. 2008. A new class of homoserine lactone quorum-sensing signals. Nature 454:595–9.

29. Lindemann A, Pessi G, Schaefer AL, Mattmann ME, Christensen QH, Kessler A, Hennecke H, Blackwell HE, Greenberg EP, Harwood CS. 2011. Isovaleryl-homoserine lactone, an unusual branched-chain quorum-sensing signal from the soybean symbiont Bradyrhizobium japonicum. Proc Natl Acad Sci USA 108:16765–70.

30. Ahlgren NA, Harwood CS, Schaefer AL, Giraud E, Greenberg EP. 2011. Aryl-homoserine lactone quorum sensing in stem-nodulating photosynthetic bradyrhizobia. 108:7183–7188.

31. Liao L, Schaefer AL, Coutinho BG, Brown PJB, Greenberg EP. 2018. An aryl-homoserine lactone quorum-sensing signal produced by a dimorphic prosthecate bacterium. 115:7587–7592.

32. Shiner EK, Rumbaugh KP, Williams SC. 2005. Interkingdom signaling: Deciphering the language of acyl homoserine lactones. FEMS Microbiol Rev 29:935–947.

33. Paggi RA, Martone CB, Fuqua C, Castro RE. 2003. Detection of quorum sensing signals in the haloalkaliphilic archaeonNatronococcus occultus. FEMS Microbiol Lett 221:49–52.

34. Zhang G, Zhang F, Ding G, Li J, Guo X, Zhu J, Zhou L, Cai S, Liu X, Luo Y, Zhang G, Shi W, Dong X. 2012. Acyl homoserine lactone-based quorum sensing in a methanogenic archaeon. ISME J 6:1336–44.

35. Charlesworth J, Beloe C, Watters C, Burns B. 2017. Quorum Sensing in Archaea: Recent Advances and Emerging Directions, p 119–132 doi:10.1007/978-3-319-65536-9_8.

36. Biswa P, Doble M. 2013. Production of acylated homoserine lactone by Gram-positive bacteria isolated from marine water. FEMS Microbiol Lett 343:34–41.

37. Bose U, Ortori CA, Sarmad S, Barrett DA, Hewavitharana AK, Hodson MP, Fuerst JA, Shaw PN, Boden R. 2017. Production of N-acyl homoserine lactones by the sponge-associated marine actinobacteria Salinispora arenicola and Salinispora pacifica. FEMS Microbiol Lett 364.

38. Ma ZP, Lao YM, Jin H, Lin GH, Cai ZH, Zhou J. 2016. Diverse Profiles of AI-1 Type Quorum Sensing Molecules in Cultivable Bacteria from the Mangrove (Kandelia obovata) Rhizosphere Environment. Front Microbiol 7:1957.

39. Charlesworth JC, Watters C, Wong HL, Visscher PT, Burns BP. 2019. Isolation of novel quorum-sensing active bacteria from microbial mats in Shark Bay Australia. FEMS Microbiol Ecol 95.

40. Mangano S, Caruso C, Michaud L, Lo Giudice A. 2018. First evidence of quorum sensing activity in bacteria associated with Antarctic sponges. Polar Biol doi:10.1007/s00300-018-2296-3.

41. Ji M, Greening C, Vanwonterghem I, Carere CR, Bay SK, Steen JA, Montgomery K, Lines T, Beardall J, van Dorst J, Snape I, Stott MB, Hugenholtz P, Ferrari BC. 2017. Atmospheric trace gases support primary production in Antarctic desert surface soil. Nature 552:400–403.

42. van Dorst J, Bissett A, Palmer AS, Brown M, Snape I, Stark JS, Raymond B, McKinlay J, Ji M, Winsley T, Ferrari BC. 2014. Community fingerprinting in a sequencing world. FEMS Microbiol Ecol 89:316–30.

43. Bay S, Ferrari B, Greening C. 2018. Life without water: How Do Bacteria Generate Biomass in Desert Ecosystems. Microbiol Aust 39:28–32.

44. Sambrook J, Russell DW. 2006. Purification of nucleic acids by extraction with phenol:chloroform. CSH Protoc 2006.

45. Weisburg WG, Barns SM, Pelletier DA, Lane DJ. 1991. 16S ribosomal DNA amplification for phylogenetic study. J Bacteriol 173:697–703.

46. McClean KH, Winson MK, Fish L, Taylor A, Chhabra SR, Camara M, Daykin M, Lamb JH, Swift S, Bycroft BW, Stewart GSAB, Williams P. 1997. Quorum sensing and Chromobacterium violaceum: exploitation of violacein production and inhibition for the detection of N-acylhomoserine lactones. 143:3703–3711.

47. Farrand SK, Qin Y, Oger P. 2002. Quorum-sensing system of Agrobacterium plasmids: Analysis and utility, p 452–484, Methods Enzymol, vol 358. Academic Press.

48. Charlesworth J, Kimyon O, Manefield M, Burns BP. 2015. Detection and characterization of N-acyl-l-homoserine lactones using GFP-based biosensors in conjunction with thin-layer chromatography. J Microbiol Methods 118:164–7.

49. Chu W, Vattem DA, Maitin V, Barnes MB, McLean RJ. 2011. Bioassays of quorum sensing compounds using Agrobacterium tumefaciens and Chromobacterium violaceum. Methods Mol Biol 692:3–19.

50. Shaw PD, Ping G, Daly SL, Cha C, Cronan JE, Rineheart KL, Farrahand SK. 1997. Detecting and characterizing N-acyl-homoserine lactone signal molecules by thin-layer chromatography. Proc Natl Acad Sci USA 94:6036–6041.

51. Chin C-S, Alexander DH, Marks P, Klammer AA, Drake J, Heiner C, Clum A, Copeland A, Huddleston J, Eichler EE, Turner SW, Korlach J. 2013. Nonhybrid, finished microbial genome assemblies from long-read SMRT sequencing data. Nat Methods 10:563.

52. Hunt M, Silva ND, Otto TD, Parkhill J, Keane JA, Harris SR. 2015. Circlator: automated circularization of genome assemblies using long sequencing reads. Genome Biology 16:294.

53. Seemann T. 2014. Prokka: rapid prokaryotic genome annotation. Bioinformatics 30:2068–2069.

54. Wu S, Zhu Z, Fu L, Niu B, Li W. 2011. WebMGA: a customizable web server for fast metagenomic sequence analysis. BMC Genomics 12:444.

55. Tatusov RL, Galperin MY, Natale DA, Koonin EV. 2000. The COG database: a tool for genome-scale analysis of protein functions and evolution. Nucleic Acids Res 28:33–36.

56. Parks DH, Imelfort M, Skennerton CT, Hugenholtz P, Tyson GW. 2015. CheckM: assessing the quality of microbial genomes recovered from isolates, single cells, and metagenomes. Genome Res 25:1043–55.

57. Jones P, Binns D, Chang H-Y, Fraser M, Li W, McAnulla C, McWilliam H, Maslen J, Mitchell A, Nuka G, Pesseat S, Quinn AF, Sangrador-Vegas A, Scheremetjew M, Yong S- Y, Lopez R, Hunter S. 2014. InterProScan 5: genome-scale protein function classification. Bioinformatics (Oxford, England) 30:1236–1240.

58. Yang J, Roy A, Zhang Y. 2012. BioLiP: a semi-manually curated database for biologically relevant ligand–protein interactions. Nucleic Acids Res 41:D1096–D1103.

59. Yang J, Roy A, Zhang Y. 2013. Protein–ligand binding site recognition using complementary binding-specific substructure comparison and sequence profile alignment. Bioinformatics 29:2588–2595.

60. Madeira F, Park Ym, Lee J, Buso N, Gur T, Madhusoodanan N, Basutkar P, Tivey ARN, Potter SC, Finn RD, Lopez R. 2019. The EMBL-EBI search and sequence analysis tools APIs in 2019. Nucleic Acids Res 47:W636–W641.

61. Dereeper A, Guignon V, Blanc G, Audic S, Buffet S, Chevenet F, Dufayard JF, Guindon S, Lefort V, Lescot M, Claverie JM, Gascuel O. 2008. Phylogeny.fr: robust phylogenetic analysis for the non-specialist. Nucleic Acids Res 36:W465–W469.

62. Consortium TU. 2014. UniProt: a hub for protein information. Nucleic Acids Res 43:D204–D212.

63. Edgar RC. 2004. MUSCLE: multiple sequence alignment with high accuracy and high throughput. Nucleic Acids Res 32:1792–1797.

64. Fierer N, Bradford MA, Jackson RB. 2007. Toward an ecological classification of soil bacteria. Ecology 88:1354–64.

65. Qin S, Li W-J, Klenk H-P, Hozzein WN, Ahmed I. 2019. Editorial: Actinobacteria in Special and Extreme Habitats: Diversity, Function Roles and Environmental Adaptations, Second Edition. 10.

66. Sun Y, Shi Y-L, Wang H, Zhang T, Yu L-Y, Sun H, Zhang Y-Q. 2018. Diversity of Bacteria and the Characteristics of Actinobacteria Community Structure in Badain Jaran Desert and Tengger Desert of China. 9.

67. Dhakal D, Pokhrel AR, Shrestha B, Sohng JK. 2017. Marine Rare Actinobacteria: Isolation, Characterization, and Strategies for Harnessing Bioactive Compounds. 8.

68. de Lima Procópio RE, da Silva IR, Martins MK, de Azevedo JL, de Araújo JM. 2012. Antibiotics produced by Streptomyces. The Brazilian Journal of Infectious Diseases 16:466–471.

69. Song L, Liu H, Wang J, Huang Y, Dai X, Han X, Zhou Y. 2015. Belliella marina sp. nov., isolated from seawater. 65:4353–4357.

70. Liu M, Qi H, Luo X, Dai J, Peng F, Fang C. 2012. Cesiribacter roseus sp. nov., a pink-pigmented bacterium isolated from desert sand. 62:96–99.

71. Gallego V, Sánchez-Porro C, García MT, Ventosa A. 2006. Roseomonas aquatica sp. nov., isolated from drinking water. 56:2291–2295.

72. Prieto-Barajas CM, Valencia-Cantero E, Santoyo G. 2018. Microbial mat ecosystems: Structure types, functional diversity, and biotechnological application. Electron J Biotechnol 31:48–56.

73. Navarro Llorens JM, Tormo A, Martínez-García E. 2010. Stationary phase in gram-negative bacteria. 34:476–495.

74. Case RJ, Labbate M, Kjelleberg S. 2008. AHL-driven quorum-sensing circuits: their frequency and function among the Proteobacteria. The Isme Journal 2:345.

75. Bassler BL. 2002. Small Talk: Cell-to-Cell Communication in Bacteria. Cell 109:421–424.

76. Lyon GJ, Novick RP. 2004. Peptide signaling in Staphylococcus aureus and other Gram-positive bacteria. Peptides 25:1389–1403.

77. Dirix G, Monsieurs P, Dombrecht B, Daniels R, Marchal K, Vanderleyden J, Michiels J. 2004. Peptide signal molecules and bacteriocins in Gram-negative bacteria: a genome-wide in silico screening for peptides containing a double-glycine leader sequence and their cognate transporters. Peptides 25:1425–1440.

78. Decho AW, Visscher PT, Ferry J, Kawaguchi T, He L, Przekop KM, Norman RS, Reid RP. 2009. Autoinducers extracted from microbial mats reveal a surprising diversity of N-acylhomoserine lactones (AHLs) and abundance changes that may relate to diel pH. Environ Microbiol 11:409–20.

79. Yates EA, Philipp B, Buckley C, Atkinson S, Chhabra SR, Sockett RE, Goldner M, Dessaux Y, Cámara M, Smith H, Williams P. 2002. N-Acylhomoserine Lactones Undergo Lactonolysis in a pH-, Temperature-, and Acyl Chain Length-Dependent Manner during Growth of Yersinia pseudotuberculosis and Pseudomonas aeruginosa. 70:5635–5646.

80. Pongsilp N, Triplett EW, Sadowsky MJ. 2005. Detection of Homoserine Lactone-Like Quorum Sensing Molecules in Bradyrhizobium Strains. Curr Microbiol 51:250–254.

81. Hanzelka BL, Stevens AM, Parsek MR, Crone TJ, Greenberg EP. 1997. Mutational analysis of the Vibrio fischeri LuxI polypeptide: critical regions of an autoinducer synthase. 179:4882–4887.

82. Hunter S, Jones P, Mitchell A, Apweiler R, Attwood TK, Bateman A, Bernard T, Binns D, Bork P, Burge S, de Castro E, Coggill P, Corbett M, Das U, Daugherty L, Duquenne L, Finn RD, Fraser M, Gough J, Haft D, Hulo N, Kahn D, Kelly E, Letunic I, Lonsdale D, Lopez R, Madera M, Maslen J, McAnulla C, McDowall J, McMenamin C, Mi H, Mutowo-Muellenet P, Mulder N, Natale D, Orengo C, Pesseat S, Punta M, Quinn AF, Rivoire C, Sangrador-Vegas A, Selengut JD, Sigrist CJA, Scheremetjew M, Tate J, Thimmajanarthanan M, Thomas PD, Wu CH, Yeats C, Yong S-Y. 2012. InterPro in 2011: new developments in the family and domain prediction database. Nucleic Acids Res 40:D306–D312.

83. Egland KA, Greenberg EP. 2001. Quorum Sensing in Vibrio fischeri: Analysis of the LuxR DNA Binding Region by Alanine-Scanning Mutagenesis. 183:382–386.

84. Voth D, J Broederdorf L, G Graham J. 2012. Bacterial Type IV Secretion Systems: Versatile Virulence Machines. Future microbiology 7:241–57.

85. Fullner KJ, Stephens KM, Nester EWJM, MGG GG. 1994. An essential virulence protein of Agrobacterium tumefaciens, VirB4, requires an intact mononucleotide binding domain to function in transfer of T-DNA. 245:704–715.

86. Patil VA, Greenberg ML. 2013. Cardiolipin-Mediated Cellular Signaling, p 195–213. In Capelluto DGS (ed), Lipid-mediated Protein Signaling doi:10.1007/978-94-007-6331-9_11. Springer Netherlands, Dordrecht.

87. Fuqua WC, Winans SC. 1994. A LuxR-LuxI type regulatory system activates Agrobacterium Ti plasmid conjugal transfer in the presence of a plant tumor metabolite. 176:2796–2806.

88. Webber MA, Piddock LJV. 2003. The importance of efflux pumps in bacterial antibiotic resistance. J Antimicrob Chemother 51:9–11.

89. Fuqua C, Greenberg EP. 2002. Listening in on bacteria: acyl-homoserine lactone signalling. Nature Reviews Molecular Cell Biology 3:685–695.

